# Algorithm for selecting potential SARS-CoV-2 dominant variants based on POS-NT frequency

**DOI:** 10.1101/2023.12.05.570216

**Authors:** Eunhee Kang, TaeJin Ahn, Taesung Park

## Abstract

COVID-19, currently prevalent worldwide, is caused by a novel coronavirus, SARS-CoV-2. Similar to other RNA viruses, SARS-CoV-2 continues to evolve through random mutations, creating numerous variants, such as Alpha, Beta, and Delta. It is, therefore, necessary to predict the mutations constituting the dominant variant before they are generated. This can be achieved by continuously monitoring the mutation trends and patterns. Hence, in the current study, we sought to design a dominant variant candidate (DVC) selection algorithm. To this end, we obtained COVID-19 sequence data from GISAID and extracted position-nucleotide (POS-NT) frequency ratio data by country and date through data preprocessing. We then defined the dominant dates for each variant in the USA and developed a frequency ratio prediction model for each POS-NT. Based on this model, we applied DVC criteria to develop the selection algorithm, verified for Delta and Omicron. Using Condition 3 as the DVC criterion, 69 and 102 DVC POS-NTs were identified for Delta and Omicron an average of 47 and 82 days before the dominant dates, respectively. Moreover, 13 and 44 Delta- and Omicron-defining POS-NTs were recognized 18 and 25 days before the dominant dates, respectively. We identified all DVC POS-NTs before the dominant dates, including soaring and gently increasing POS-NTs. Considering that we successfully defined all POS-NT mutations for Delta and Omicron, the DVC algorithm may represent a valuable tool for providing early predictions regarding future variants, helping improve global health.

**Author Summary:** 

## Introduction

The recent outbreak of COVID-19, caused by the novel SARS-CoV-2 virus, has severely affected global society. Continuous mutations in the genome generate new variants that enable the virus to thwart disease control measures. Next-generation sequencing technology is widely employed to characterize the genetic SARS-CoV-2 variants. Owing to the contributions of many researchers, SARS-CoV-2 genomic data has been collected from infected individuals worldwide. GISAID is a database that stores and provides sequenced SARS-COV-2 genomes along with basic metadata, including the sequencing date and location. GISAID also presents the status of SARS-CoV-2 variant spread visually in a geographical and time-dependent manner. In particular, predicting the emergence of a novel variant is important to identify new potential outbreaks capable of evading the current diagnostic and vaccine strategies.

In this study, we provide a prediction model that estimates whether a single SARS-COV-2 mutation is a prominent factor in determining disease severity in infected patients. This functionality is helpful in disease control in several aspects. First, single mutations may be associated with known clinical characteristics, such as symptom severity, incubation period, and morbidity rate. Second, mutations in PCR primer binding regions can be used to estimate if an infectious virus evades diagnostic methods. Third, single mutations help assess the vaccination efficacy of the designed epitope.

### SNPs defining Delta and Omicron variants

Based on the WHO nomenclature system (GISAID, Pango lineage, Nextstrain clade): Alpha (GRY, B.1.1.7, 20I (V1)), Beta (GH/501Y. V2, B.1.351, 20H (V2)), Gamma (GR/501Y.V3, P.1, 20J(V3)), Delta (G/478K.V1, B.1.617.2, 21A-21I-21J), and Omicron (GR/484A, B.1.1.529, 21K-21L-21M-22A-22B-22C-22D) (WHO: https://www.who.int/activities/tracking-SARS-CoV-2-variants) SARS-CoV-2 strains have arisen due to mutations in the genomic sequence. The SARS-CoV-2 genome comprises 29,903 nucleotides, encoding 12 proteins (ORF1a/1ab, S, ORF3a, ORF3b, E, M, ORF6, ORF7a, ORF7b, ORF8, and ORF10*). These mutations are caused by single nucleotide changes, i.e., replacement, insertion, or deletion, leading to changes in the amino acid sequence (Fig 1; GISAID: https://gisaid.org/). Fig 1 presents a genome sequence map of SARS-Cov-2 and the major mutational positions of several variants. In this study, we attempted to predict the Delta and Omicron variants, the most recent dominant SARS-CoV-2 variants. According to Pango (Pango cov-lineages: https://cov-lineages.org/), the Delta and Omicron variants have 13 and 47 defining SNPs, respectively (Tables 1 and 2).

**Fig 1.**
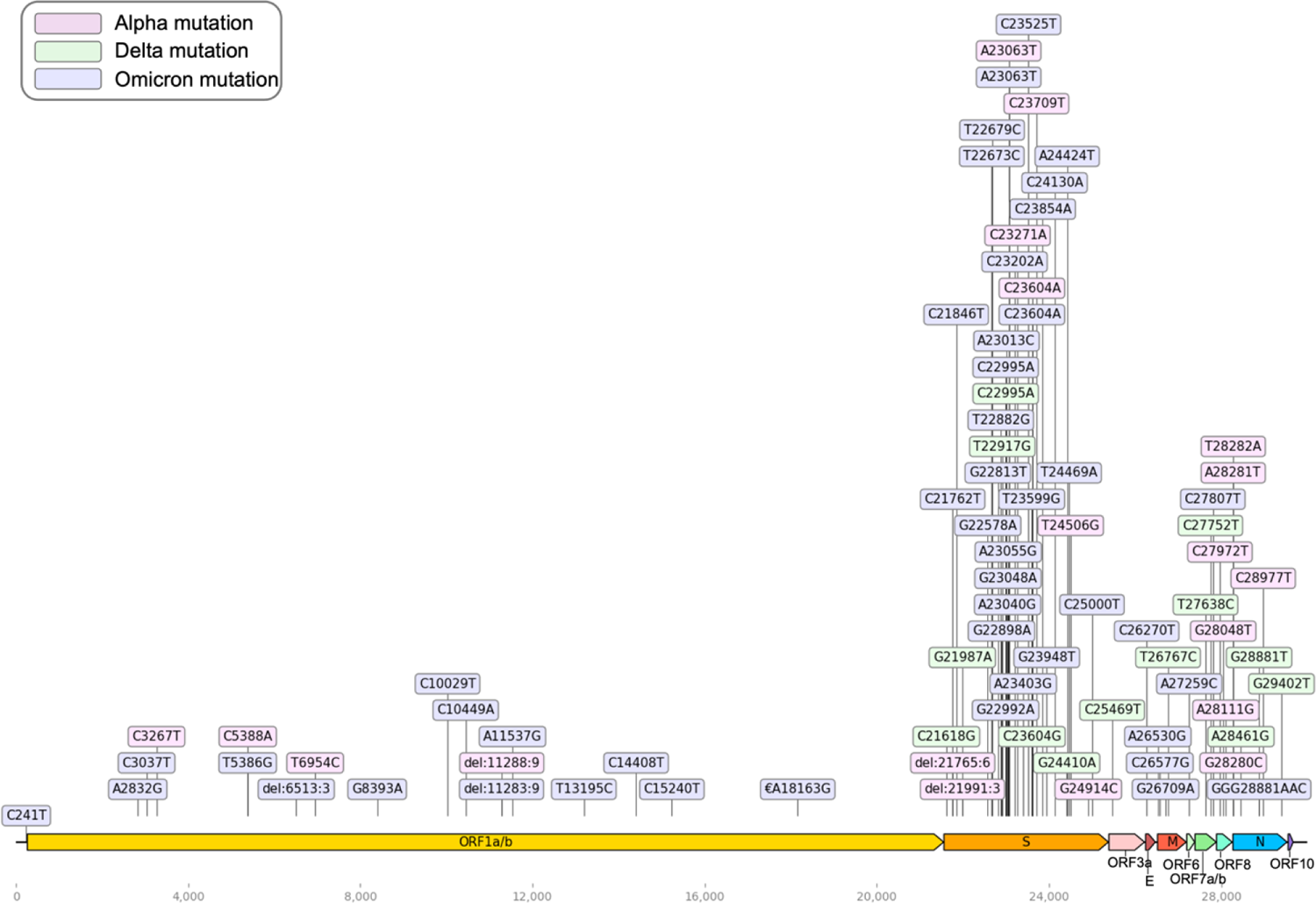
SARS-CoV-2 genome sequence map. The 29,903 nucleotide positions are shown in the context of the 12 encoded proteins. The main mutations of each dominant variant are shown; each color indicates each mutation. Pink: Alpha mutation, Green: Delta mutation, Blue: Omicron mutation.

**Table 1.** Delta defining position-nucleotide list.

|  | **Amino acid** | **Nucleotide** |
| --- | --- | --- |
| **1** | T19R | C21618G |
| **2** | T478K | C22995A |
| **3** | L452R | T22917G |
| **4** | D950N | G24410A |
| **5** | P681R | C23604G |
| **6** | D377Y | G21987A |
| **7** | S26L | T26767C |
| **8** | I82T | T27638C |
| **9** | V82A | C27752T |
| **10** | R203M | C25469T |
| **11** | T120I | G29402T |
| **12** | G142D | A28461G |
| **13** | D63G | G28881T |

**Table 2.** Omicron defining position-nucleotide list.

|  | **Amino acid** | **Nucleotide** |
| --- | --- | --- |
| **1** | **6513>3** |  |
| **2** | **11283>9** |  |
| **3** | C241T | C241T |
| **4** | K856R | A2832G |
| **5** | C3037T | C3037T |
| **6** | T5386G | T5386G |
| **7** | A2710T | G8393A |
| **8** | T3255I | C10029T |
| **9** | P3395H | C10449A |
| **10** | I3758V | A11537G |
| **11** | T13195C | T13195C |
| **12** | P314L | C14408T |
| **13** | C15240T | C15240T |
| **14** | I1566V | A18163G |
| **15** | A67V | C21762T |
| 16 | T95I | C21846T |
| 17 | G339D | G22578A |
| 18 | S371L | T22673C |
| 19 | S373P | T22679C |
| 20 | K417N | G22813T |
| 21 | N440K | T22882G |
| 22 | G446S | G22898A |
| 23 | S477N | G22992A |
| 24 | T478K | C22995A |
| 25 | E484A | A23013C |
| 26 | Q493R | A23040G |
| 27 | G496S | G23048A |
| 28 | Q498R | A23055G |
| 29 | N501Y | A23063T |
| 30 | T547K | C23202A |
| 31 | D614G | A23403G |
| 32 | H655Y | C23525T |
| 33 | N679K | T23599G |
| 34 | P681H | C23604A |
| 35 | N764K | C23854A |
| 36 | D796Y | G23948T |
| 37 | N856K | C24130A |
| 38 | Q954H | A24424T |
| 39 | N969K | T24469A |
| 40 | C25000T | C25000T |
| 41 | T9I | C26270T |
| 42 | D3G | A26530G |
| 43 | Q19E | C26577G |
| 44 | A63T | G26709A |
| 45 | A27259C | A27259C |
| 46 | C27807T | C27807T |
| 47 | **RG203KR** | **GGG28881AAC** |

### COVID-19 sequence data from GISAID

The Global Initiative for Sharing All Influenza Data (GISAID) is an organization that provides a database of nucleotide sequence information and related epidemiological information of all influenza viruses and COVID-19-causing coronaviruses. GISAID provides multiple SARS-CoV-2 sequence data analyses collected worldwide, as well as sequence alignments, diagnostic primer and probe coordinates, 3D protein models, drug targets, and phylogenetic trees. In this study, global SARS-CoV-2 sequence data were obtained from GISAID on February 22, 2022; 8,474,962 sequence data were obtained from December 1, 2019, to February 22, 2022 (GISAID: https://gisaid.org/).

### Data preprocessing and formatting

COVID-19 sequence data obtained in a FASTA file format from GISAID were converted from a multiline to a single-line format; only complete sequences corresponding to > 29,000 bp were extracted. We secured sequence data by country and date through the GISAID unique ID, country, collection date, and sequence information in the header of the sequence data. In this study, the USA (2,702,068), UK (1,936,958), and Germany (415,309), with the most sequencing data, in addition to data from Korea, were analyzed. The sequence data obtained by country were mapped to the original sequence (NC_045512) to obtain a sequence alignment/map (SAM) file. The binary alignment/map (BAM) file was then converted to a binary format using SAMtools to reduce the file size. From the generated BAM file, sequencing reads were synthesized for each position of the original sequence to determine whether bases differed from the original sequencing data; the mutation data was extracted in a variant call format. The obtained data were used to extract information on the number of mutations and mutation frequency ratio information for each position in the sequence and to confirm the mutation trend by securing the frequency ratio data by country, date, and POS-NTs (total 29,903 positions × 4 nucleotides × 4 countries; Fig. 2).

**Fig 2.**
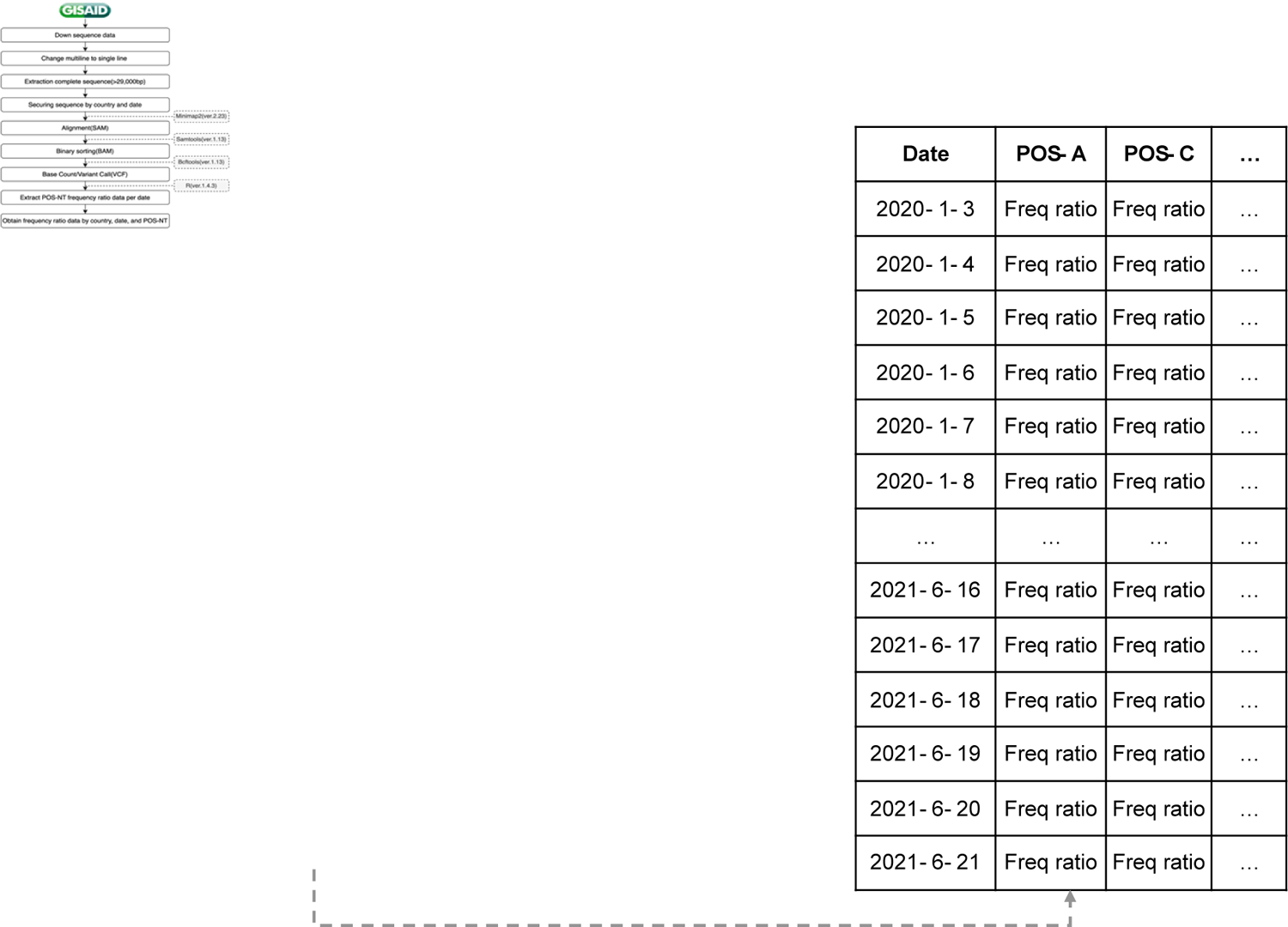
Frequency ratio data acquisition process by country, date, and POS-NT.

### Frequency (freq)

As it is overly computationally intensive to check the trend for all mutations in SARS-COV-2 (i.e., a combination of 29,903 positions and three SNP mutations), each nucleotide position was subjected to additional preprocessing to remove positions where mutations did not occur (reference frequency = 100), those with no information on the time point of the dominant variant, and positions where the change in reference allele frequency was < 10%. Next, we created continuous data for date information for which frequency ratio information did not exist, and the position where the total data date was < 50 removed. Subsequently, cubic spline interpolation was used to fill in the data for which the frequency ratio information did not exist. We removed the reference allele from the four nucleotides as we were interested in mutations. An additional preprocessing step is shown in Fig 3.

**Fig 3.**
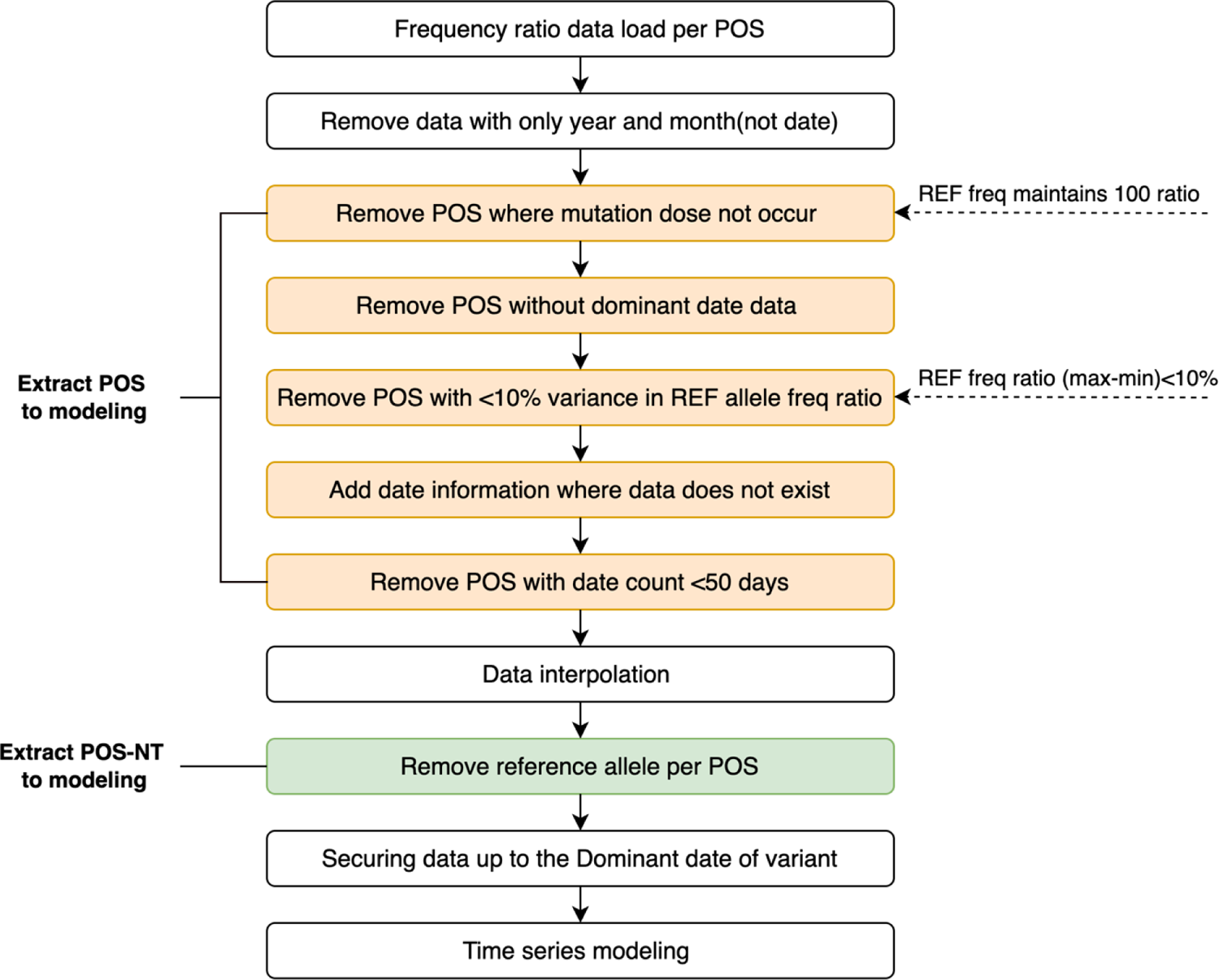
Additional preprocessing step to select POS-NTs for modeling.

### DVC selection for the prediction model

We attempted to predict the mutations comprising the dominant variants by analyzing and predicting the Delta and Omicron variants. To confirm the trend of a specific POS-NT, a dominant variant selection time point is required. Moreover, we aimed to confirm whether the developed algorithm can identify all mutations constituting the Delta and Omicron variants at the dominant variant time point after determining the DVC POS-NT until the variant became dominant. Therefore, we attempted to define the dominant variant time-point (i.e., dominant date) for Delta and Omicron. We defined the strains that accounted for > 50% of all new COVID-19 cases as the dominant variants. However, there was no information on the strain and lineage of the sequences in the data provided by GISAID. Therefore, we proceeded with the lineage analysis provided by Pangolin, assigned a strain label, including Delta and Omicron, for each sequence, and secured the daily frequency ratio data of the strain. The strain that accounted for more than 50% of all new COVID-19 cases was defined as the dominant variant, and the corresponding time point was defined as the dominant date. Fig 4 illustrates the scheme determining the dominant date for each country and its variants. The dominant date was used as the time point for selecting the dominant variant using this algorithm and as a criterion for learning and prediction date windows for each variant. For the analysis and prediction of Delta mutations, the extracted data, including the alpha-dominant to the delta-dominant dates, were employed for each POS-NT. The analysis and prediction of Omicron mutation data from the Delta dominant to Omicron dominant date.

**Fig 4.**
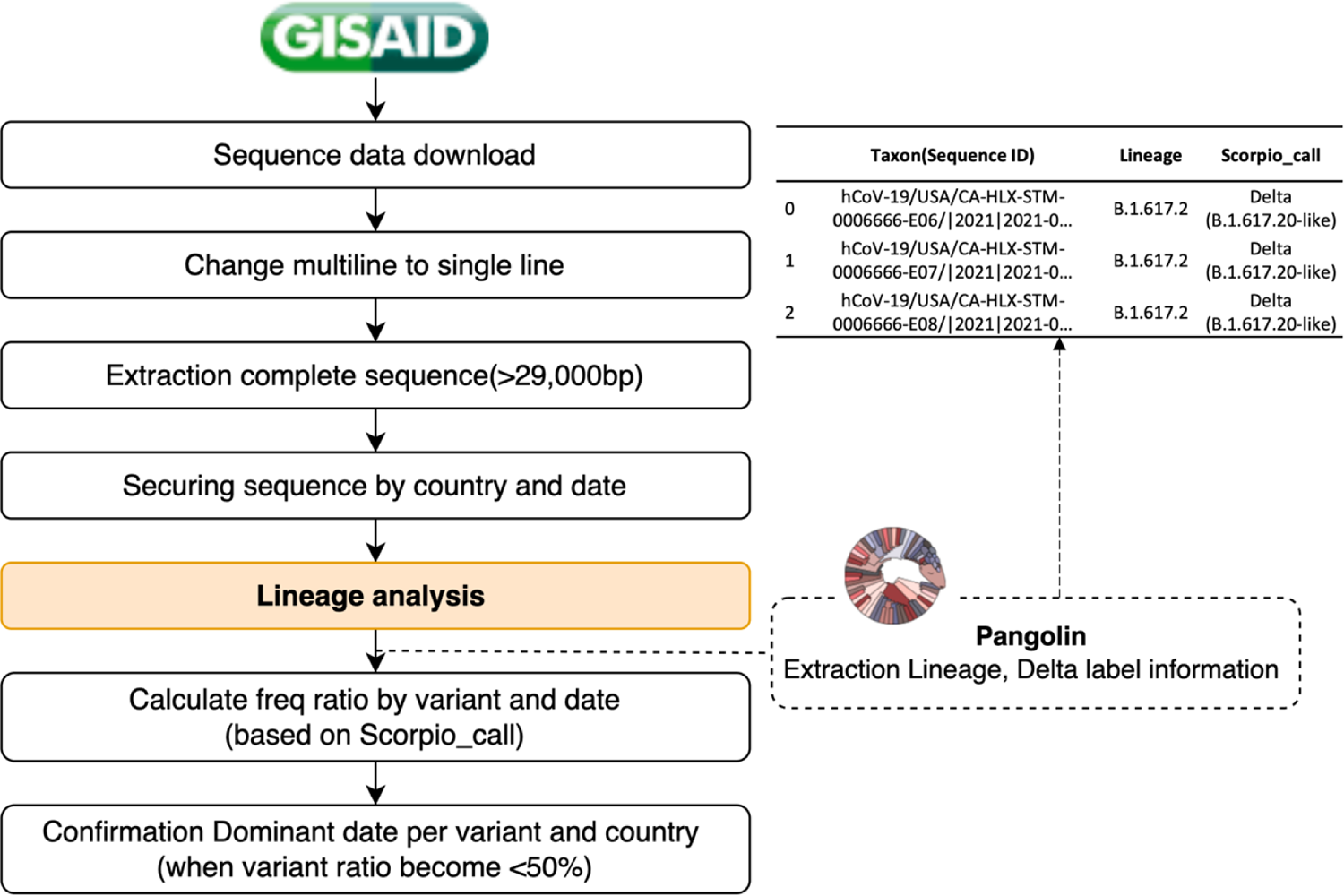
Dominant variant time point (dominant date definition process). After detecting the sequence data for each country, the strain and lineage information for each sequence was allocated through lineage analysis provided by Pangolin. After securing daily rate data for each strain, those accounting for > 50% of all new COVID-19 cases were defined as dominant, and the corresponding time point was defined as the dominant variant time point (dominant date). The dominant date was used as the dominant variant selection time point and as a criterion for learning and prediction date windows for each variant in the algorithm.

### POS-NT frequency ratio prediction model

A POS-NT frequency ratio prediction model was developed to confirm the trend in each POS-NT frequency ratio. Gaussian Process Regression (GPR) is a new powerful Bayesian-based, non-parametric kernel-based probabilistic model for regression problems applied in exploration and utilization scenarios. It predicts the output of a new test set considering the novel input vectors of the test and training sets [1–3]. The most prominent advantage of GPRs is their ability to obtain the forecast uncertainty with the forecast value. In addition, GPR boasts computational efficiency and high accuracy, and is suitable for other time series forecasting, such as weather forecasting^4^. Recently, the GPR model was used widely in predicting COVID-19 spread and deaths, exhibiting improved performance compared with other models [3,5–7]. The following four patterns were identified for Delta and Omicron variant-defining mutations: (1) a gentle increase from a ratio value of 0 (Fig 5a); (2) high-frequency ratio values are consistently present (Fig 5b); (3) a gentle increase through the dominant date, with a high-frequency ratio, of the previous variant (Fig 5c); (4) soaring pattern (Fig 5d). To identify the trend of a gently increasing pattern and soaring pattern, it was necessary to select optimal training and prediction dates. Therefore, to learn the trend of the soaring pattern, we sought to apply the latest information to predict the future and modeled each learning and prediction combination until the dominant date for each variant (i.e., learn for 10 and 20 days and predict 3, 5, 8, and 10 days later; Fig 6a). In the case of the Delta mutation, data from the Alpha-dominant to Delta-dominant dates were employed for the analysis window based on the variant. In the case of the Omicron mutation, data from the Delta-dominant date to the Omicron-dominant date were modeled (Fig 6b).

**Fig 5.**
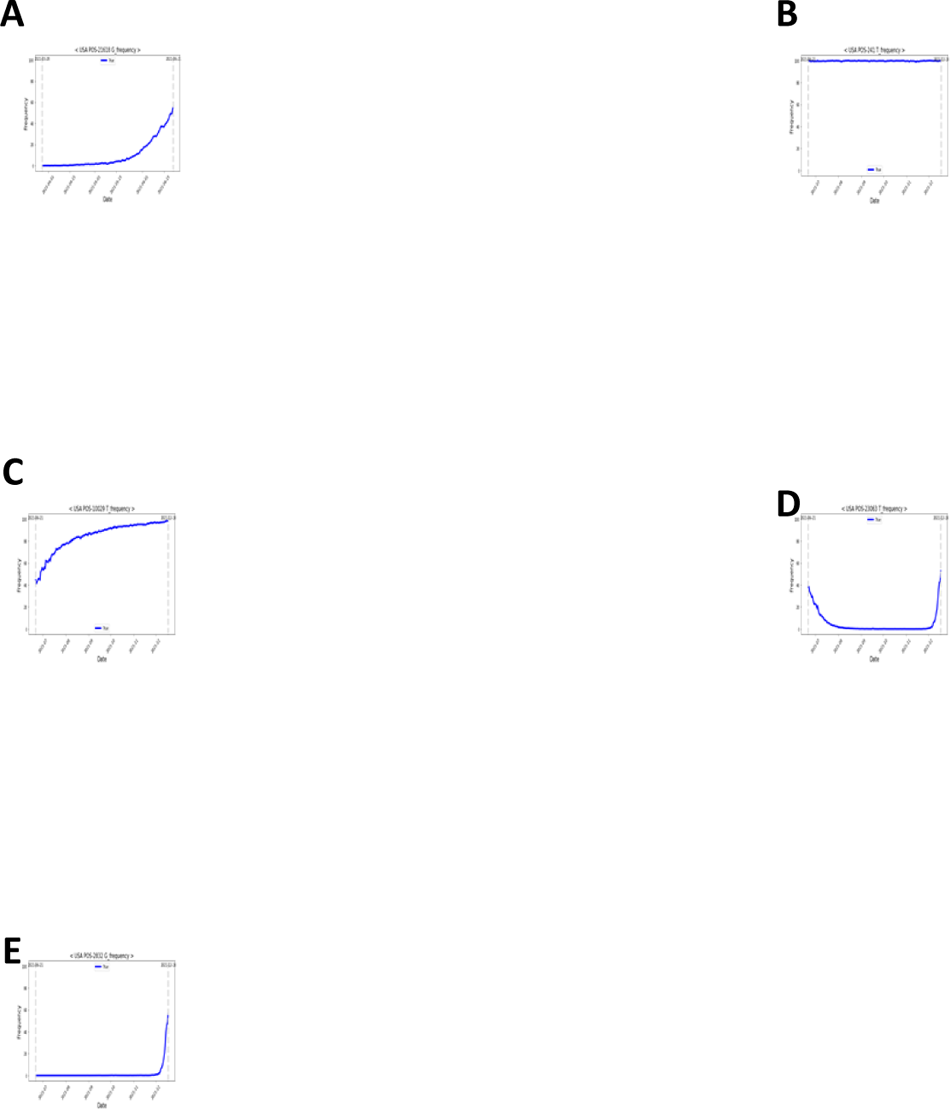
Time-dependent patterns of delta- and omicron-defining mutations. (**a**) Gentle increase from a ratio value of 0; (**b**) high-frequency ratio values are consistently present; (**c**) gentle increase through the dominant date (with a high-frequency ratio) of the previous variant; (**d**) soaring pattern. Fig **5a** shows the pattern of the Delta variant. Fig **5b–e** show the patterns of the Omicron variant.

**Fig 6.**
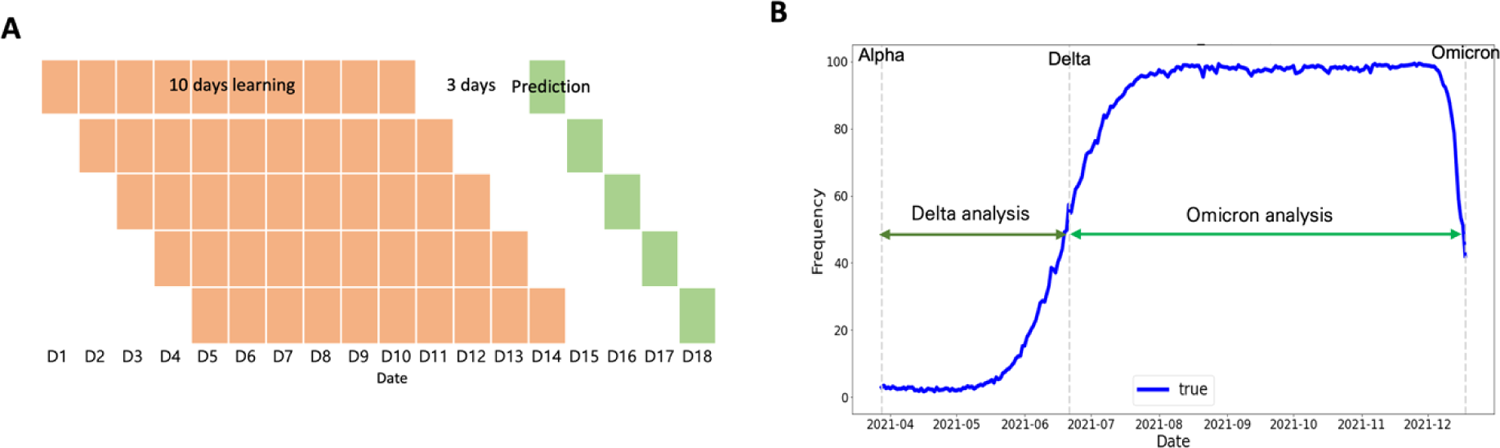
Learning and prediction window selection. (**a**) Example training and prediction window for one POS-NT (training for 10 days and predicting after 3 days) and (**b**) Delta and Omicron analysis date time.

### DVC selection algorithm

In this study, based on the frequency ratio prediction model for each POS-NT, a dominant variant candidate selection algorithm (DVC selection algorithm) was developed by applying the dominant variant candidate criteria (DVC criteria; Fig 7). We then determined whether all POS-NTs met the DVC criteria for each prediction time point; upon failing to meet the DVC criteria, the corresponding POS-NT was reanalyzed the next day. If it met the DVC criteria at that time point, the corresponding POS-NT was classified as DVC POS-NT. The DVC POS-NT was identified up to the dominant date of the variant, and then the identified DVC POS-NT was compared with the actual variant definition POS-NT list. Eight conditions were simulated to select the optimal DVC criteria based on the criteria for outliers in which the frequency of the DVC increased the next day (i.e., Criterion 2), and the measured value was higher than the predicted value (i.e., Criterion 4; Table 3). The DVC criteria defined the corresponding POS-NT as DVC POS-NT when all four detailed criteria were satisfied.

**Fig 7.**
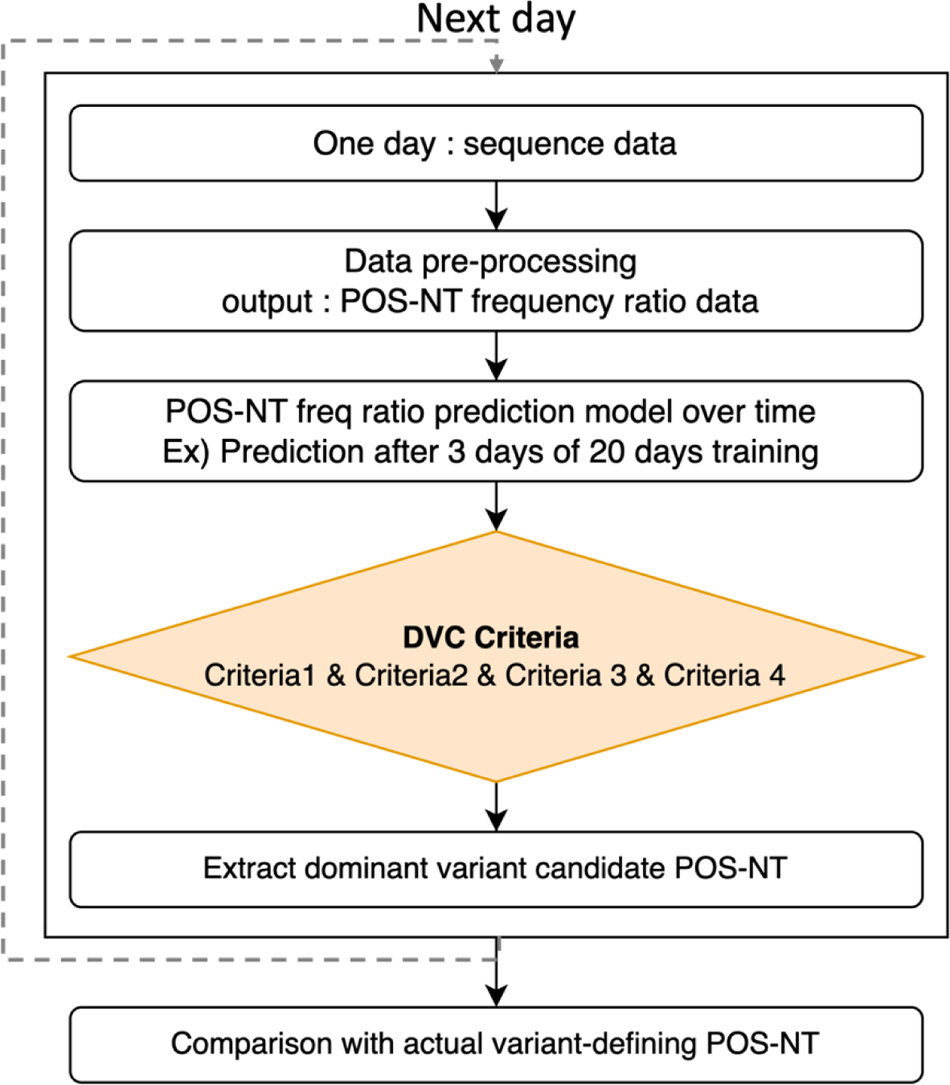
DVC POS-NT Selection Algorithm and Combined DVC Criteria. When POS-NT ratio data at a specific point occur, predictions for the future take place. If the DVC criteria were met, the corresponding POS-NT was identified as the DVC POS-NT. If it does not meet the DVC criteria at that time point, the POS-NT moves to the next time point, and the analysis continues. Criteria 1: number of days in which all dominant variant candidate criteria were satisfied; Criterion 2: whether the observed frequency ratio increased the next day compared with the previous day; Criterion 3: threshold of the predicted frequency ratio; Criterion 4: Observed value greater than the predicted value.

**Table 3.**
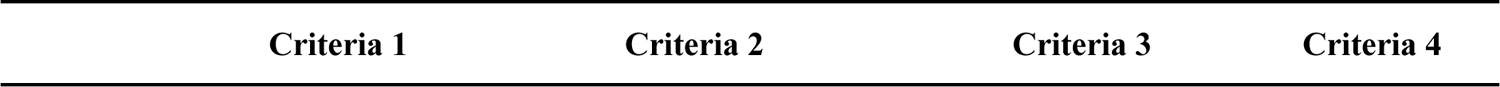
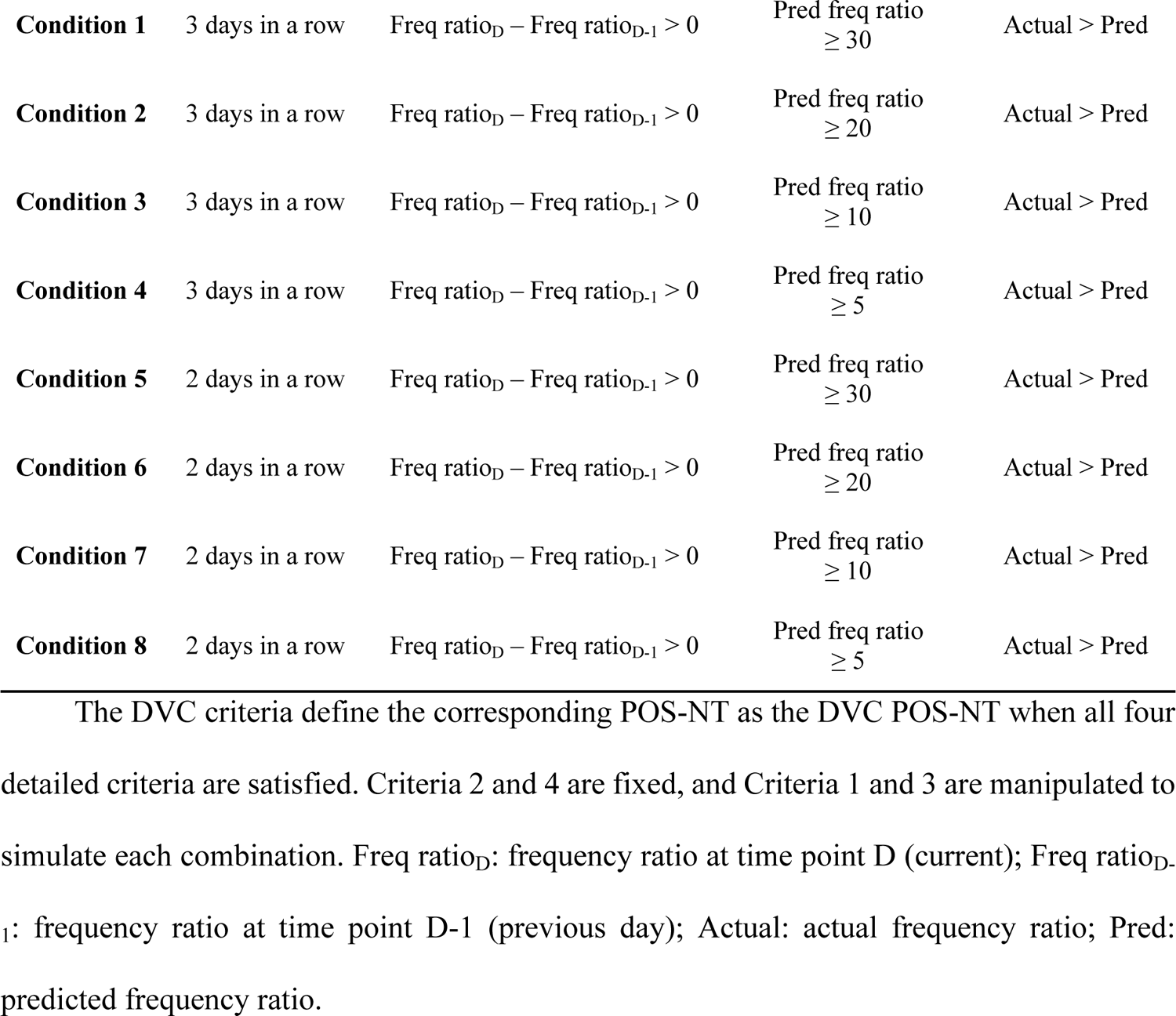
Eight DVC criteria combinations.

| Criteria 1 | Criteria 2 | Criteria 3 | Criteria 4 |
| --- | --- | --- | --- |

|  |  |  |  |  |
| --- | --- | --- | --- | --- |
| **Condition 1** | 3 days in a row | $\text{Freq ratio}_D - \text{Freq ratio}_{D-1} > 0$ | $\text{Pred freq ratio} \geq 30$ | Actual > Pred |
| **Condition 2** | 3 days in a row | $\text{Freq ratio}_D - \text{Freq ratio}_{D-1} > 0$ | $\text{Pred freq ratio} \geq 20$ | Actual > Pred |
| **Condition 3** | 3 days in a row | $\text{Freq ratio}_D - \text{Freq ratio}_{D-1} > 0$ | $\text{Pred freq ratio} \geq 10$ | Actual > Pred |
| **Condition 4** | 3 days in a row | $\text{Freq ratio}_D - \text{Freq ratio}_{D-1} > 0$ | $\text{Pred freq ratio} \geq 5$ | Actual > Pred |
| **Condition 5** | 2 days in a row | $\text{Freq ratio}_D - \text{Freq ratio}_{D-1} > 0$ | $\text{Pred freq ratio} \geq 30$ | Actual > Pred |
| **Condition 6** | 2 days in a row | $\text{Freq ratio}_D - \text{Freq ratio}_{D-1} > 0$ | $\text{Pred freq ratio} \geq 20$ | Actual > Pred |
| **Condition 7** | 2 days in a row | $\text{Freq ratio}_D - \text{Freq ratio}_{D-1} > 0$ | $\text{Pred freq ratio} \geq 10$ | Actual > Pred |
| **Condition 8** | 2 days in a row | $\text{Freq ratio}_D - \text{Freq ratio}_{D-1} > 0$ | $\text{Pred freq ratio} \geq 5$ | Actual > Pred |

The DVC criteria define the corresponding POS-NT as the DVC POS-NT when all four detailed criteria are satisfied. Criteria 2 and 4 are fixed, and Criteria 1 and 3 are manipulated to simulate each combination. Freq ratio_D_: frequency ratio at time point D (current); Freq ratio_D-1_: frequency ratio at time point D-1 (previous day); Actual: actual frequency ratio; Pred: predicted frequency ratio.

A visual summary of the methodology is shown in Fig 8. The SARS-CoV-2 sequence data from GISAID were formalized to secure frequency ratio information for each POS-NT by country and date. A time-series forecasting model was developed using the time-series frequency ratio data obtained using POS-NT. Over time, learning and prediction progressed to the dominant date for each variant. For each prediction date, the DVC POS-NT selection algorithm was applied to all POS-NTs to secure DVC POS-NT for each variant. When all DVC POS-NTs were selected until the dominant date for each variant, they were compared with the actual variant-defining POS-NT to determine the number of days preceding the dominant date of the average number of variant-defining POS-NTs. We also compared the prediction results with the actual variant-defining POS-NT, to determine how many variant-defining POS-NTs could be identified, on average, how many days ago.

**Fig 8.**
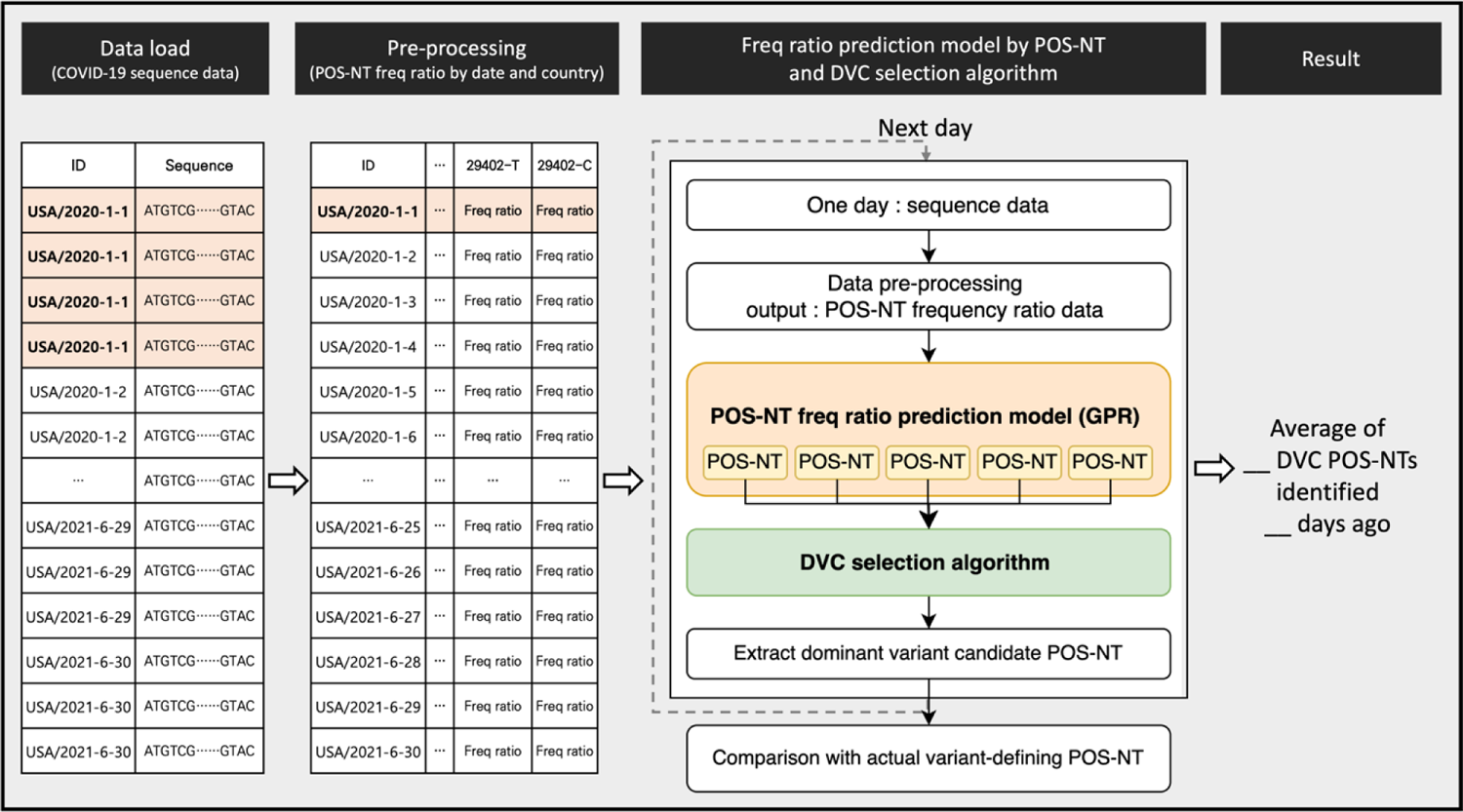
Visual summary of the methodology.

### Confirmation metric for the results

The following four metrics were used to confirm the results: (1) number of DVC POS-NTs identified by the algorithm developed in this study, i.e., candidate count; (2) average number of days for identification; (3) number of POS-NTs corresponding to the POS-NTs that define the actual variant (candidate∩actual) among the identified DVC POS-NTs; (4) ratio of the number of POS-NTs corresponding to the actual variant-defining POS-NTs among the identified DVC POS-NTs (Eq. (1)). Upon identifying all actual variant-defining POS-NT, the Candidate∩Actual value will be incremented. The algorithm can sensitively identify the DVC POS-NT as the ratio value increases.

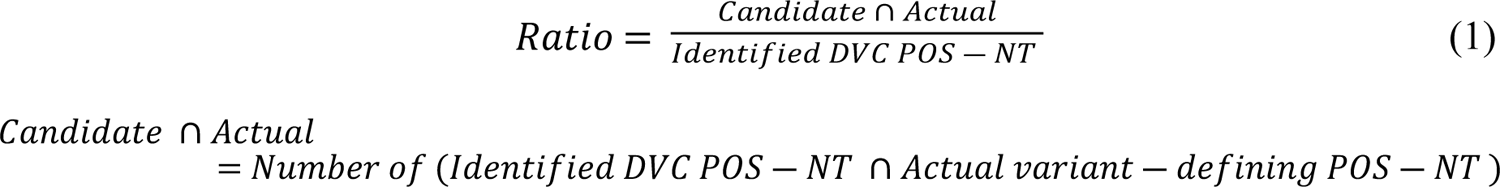

### POS-NT frequency ratio data for modeling

The numbers of nucleotide positions for the final model and the total number of models (POS-NT) are listed in Table 4. In this study, the USA data were analyzed for the first time. Prediction modeling was performed with the frequency ratio data for 6951 POS-NTs of Delta and 6990 of Omicron variants.

**Table 4.** Number of nucleotides removed during the preprocessing process and the number of POS-NTs to be modeled.

| **Table 4-1:** Number of positions removed during the preprocessing process |  |  |  |  |  |  |  |  |
| --- | --- | --- | --- | --- | --- | --- | --- | --- |
|  | Delta |  |  |  | Omicron |  |  |  |
|  | USA | UK | Korea | Germany | USA | UK | Korea | Germany |
| POS where mutation do not occur | 193 | 2399 | 15019 | 2375 | 193 | 2399 | 15019 | 2374 |
| POS without dominant date data | 1008 | 3478 | 6063 | 2292 | 99 | 327 | 274 | 240 |
| POS with < 10% variance in REF allele freq ratio | 26382 | 19962 | 4035 | 8461 | 27281 | 23110 | 9730 | 10507 |
| POS with date count < 50 days | 3 | 0 | 89 | 1 | 0 | 0 | 22 | 0 |
| **Table 4-2:** The number of POS-NTs to be modeled (the number of total models) |  |  |  |  |  |  |  |  |
|  | Delta |  |  |  | Omicron |  |  |  |
|  | USA | UK | Korea | Germany | USA | UK | Korea | Germany |
| Number of<br>POS for<br>modeling | 2317 | 4064 | 4697 | 16773 | 2330 | 4067 | 4858 | 16782 |
| Number of<br>POS-NTs for<br>modeling | 6951 | 12192 | 14091 | 50319 | 6990 | 12201 | 14574 | 50346 |

Position (POS), Frequency (freq).

### Dominant date by country and variant

In the USA, the Delta and Omicron variants were confirmed as the dominant variants on June 21 and December 18, 2021, respectively (Fig 9a). In the UK, Delta emerged as the dominant variant on May 15, 2021, and Omicron on December 14, 2021 (Fig 9b), and in Korea, Delta emerged as the dominant variant on July 4, 2021, and Omicron on January 5, 2022 (Fig 9c). In Germany, the Delta mutation was defined as the dominant variant on June 13, 2021, while the Omicron mutation accounted for > 50% of new COVID-19 cases on December 28, 2021 (Fig 9d). We used the dominant date as the dominant variant selection time point for this algorithm and as the criterion for the learning and prediction date windows for each variant.

**Fig 9.**
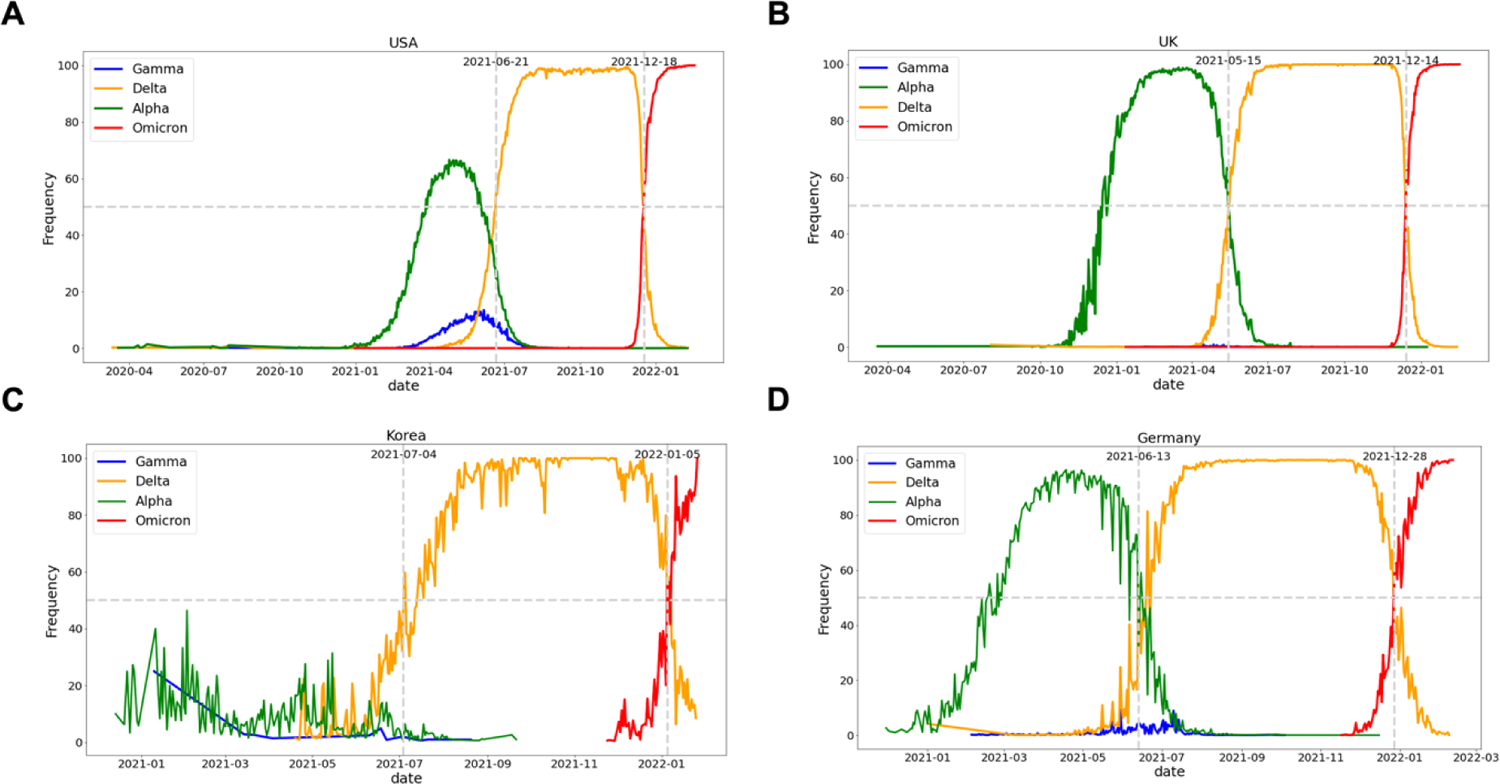
Definition of dominant dates for Delta and Omicron by country. (**a)** In the USA, Delta became the dominant variant on June 21, 2021, and Omicron on December 18, 2021, (**b)** in the UK, Delta became the dominant variant on May 15, 2021, and Omicron on December 14, 2021, (**c)** in Korea, Delta became the dominant variant on July 4, 2021, and Omicron on January 5, 2022, and (**d**) in Germany, Delta was defined as the dominant variant on June 13, 2021, and Omicron accounted for more than 50% of all new COVID-19 cases on December 28, 2021.

### POS-NT frequency ratio prediction model

The prediction results for each learning and prediction date combination, i.e., 10- and 20- day training and prediction after 3, 5, 8, and 10 days, were confirmed. Figs 10 and 11 show the Delta and Omicron predictions for a model trained for 20 days and predicted three days after the learning period. The results for the learning and prediction for other combinations are shown in S1 Fig 1–14. It was confirmed that as the number of forecast days decreased, the forecast trend improved (after 3 days > after 5 days > after 8 days > after 10 days).

**Fig 10.**
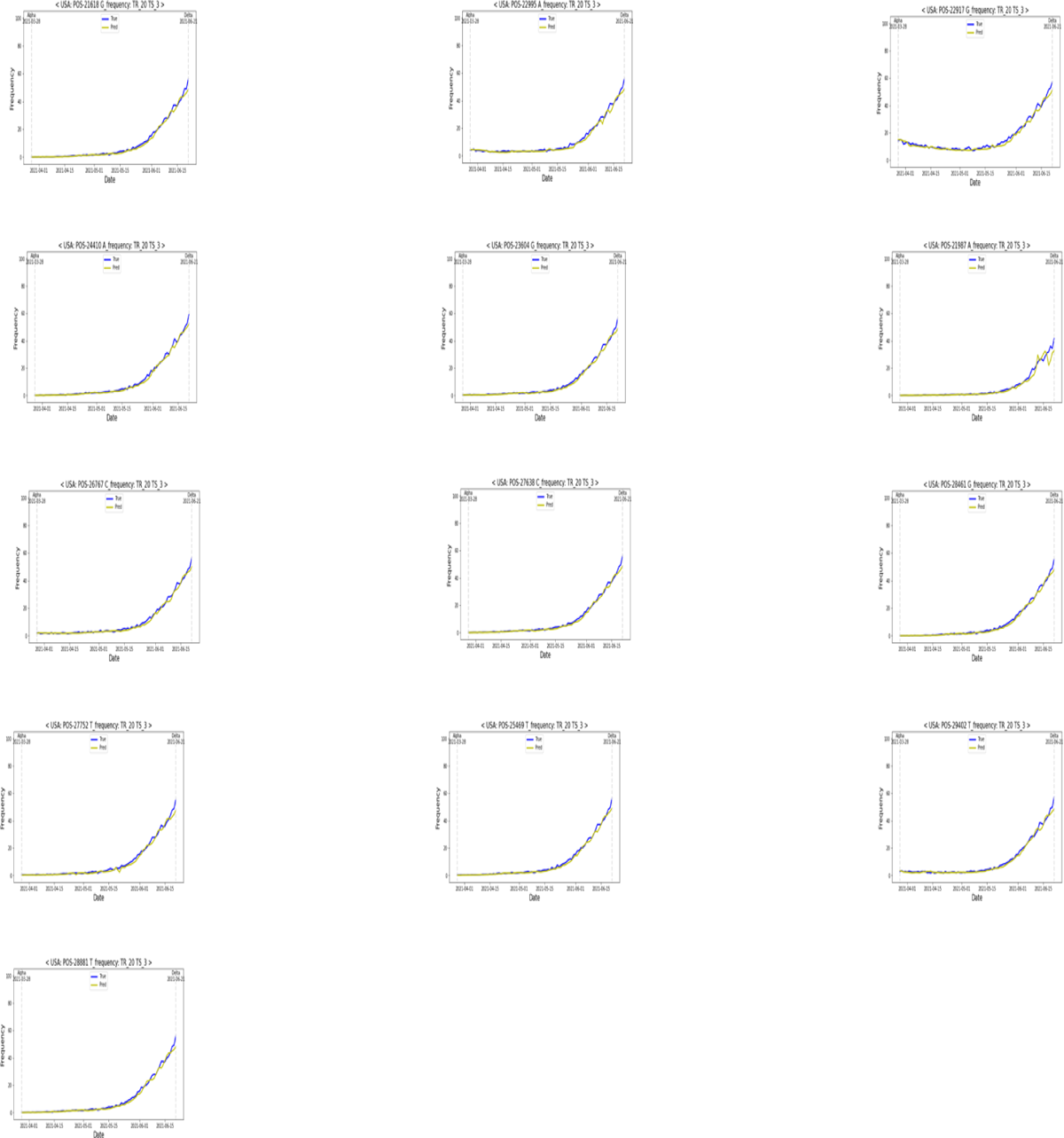
Delta: Results of learning for 20 days and predicting 3 days later. TR: learning dates (training dates), TS: test dates.

**Fig. 11.**
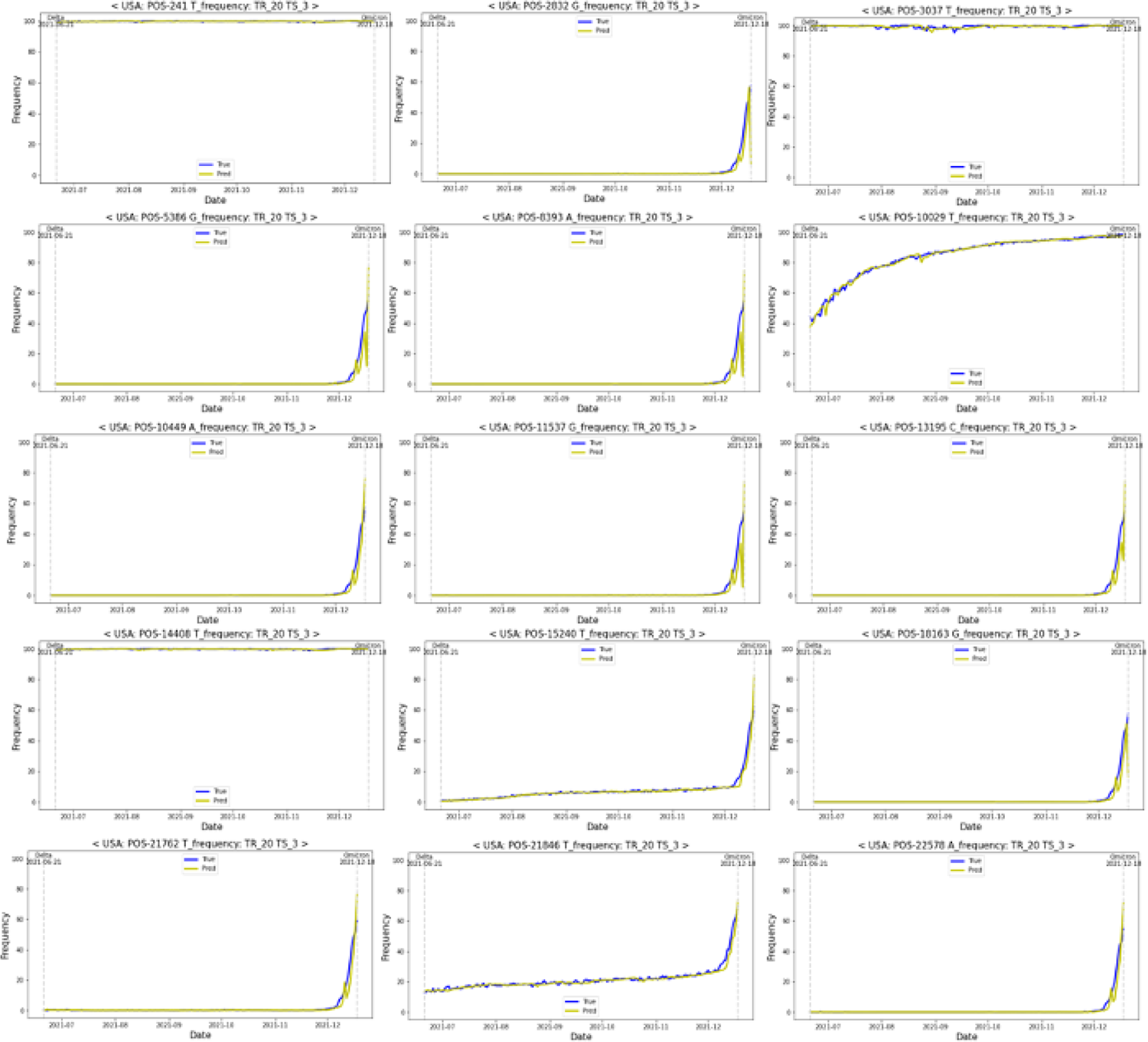
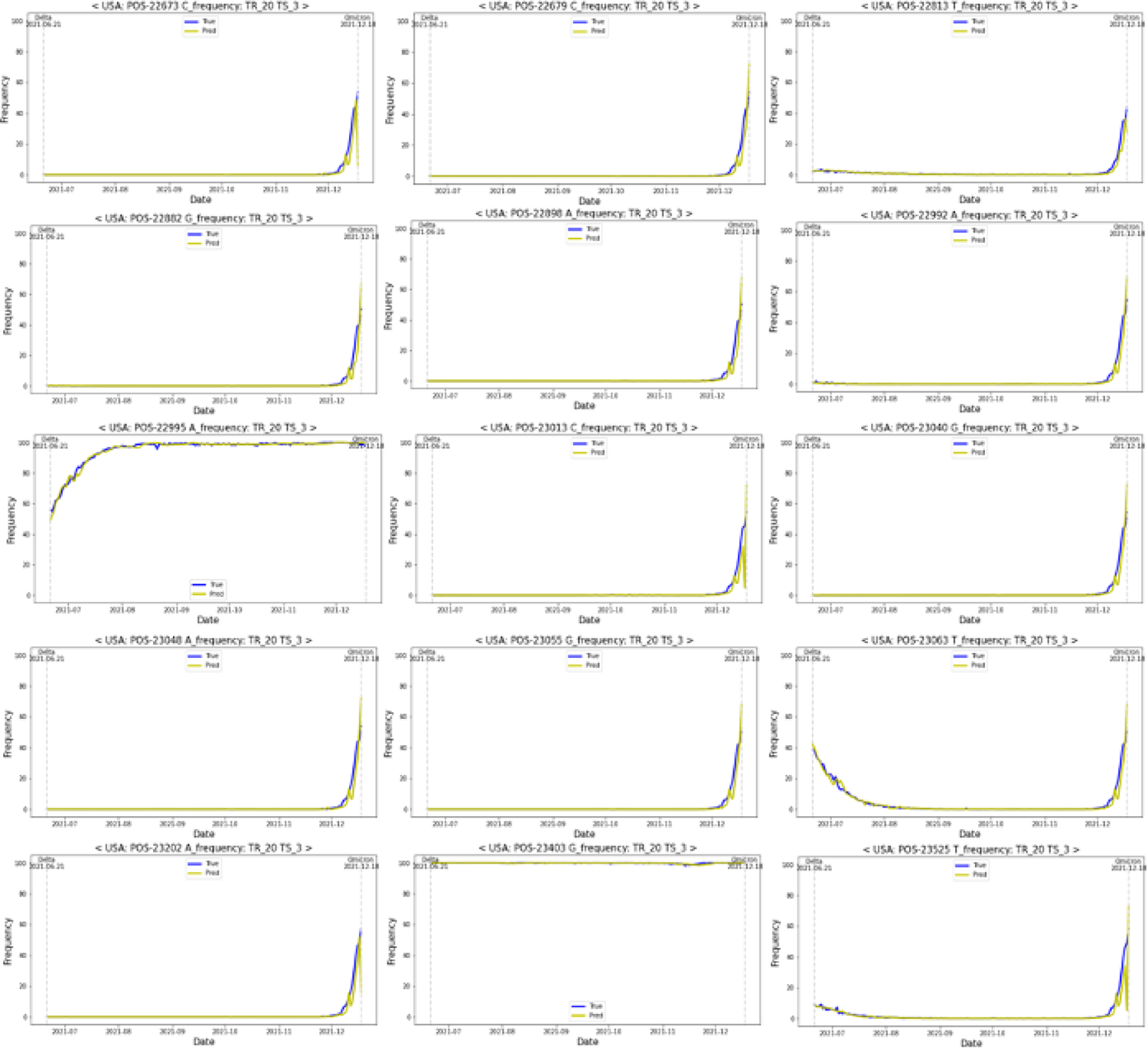
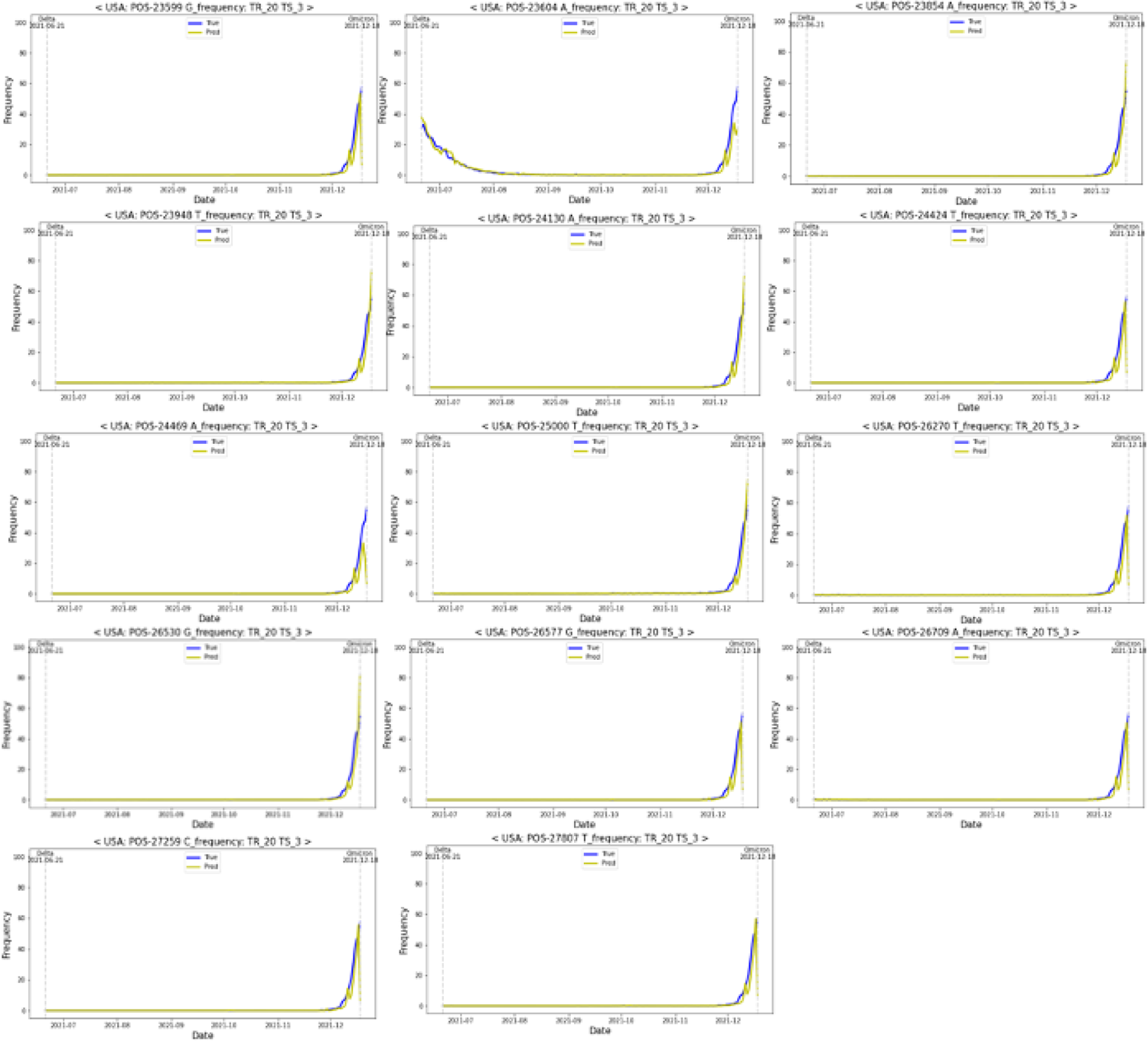
Omicron: Results of learning for 20 days and predicting 3 days later. TR: learning dates (training dates), TS: test dates.

### POS-NT identification with the algorithm

Based on the developed frequency ratio prediction model for each POS-NT, eight combinations of DVC criteria were applied to identify DVC POS-NT until the dominant date of each variant and were compared with the actual variant-defining POS-NT. In addition, the number of days ago, on average, that the POS-NT was identified as a DVC POS-NT and the ratio of the identified POS-NT corresponding to the actual variant-defining POS-NT to the identified DVC POS-NT was confirmed (Eq. (1)). S1 Table provides the learning dates, prediction dates, number of POS-NTs recognized as DVC POS-NTs by condition and the average number of days for identifying Delta mutation for all combinations of learning and prediction dates and the eight DVC conditions. S2 Table shows Delta-like information for Omicron.

The optimal DVC criterion was specified when two conditions were satisfied: (1) identify all variant-defining POS-NTs in Delta and Omicron, and (2) have the highest ratio (Eq. (1)). In the case of Delta mutation, all Delta-defined POS-NTs were identified in 39 model-specific and DVC criteria combinations and showed the highest ratio values in prediction using Condition 2, 3 days after the 20-day learning period. In the case of the Omicron mutation, all Omicron-defined POS-NTs were identified in 11 model-specific and DVC criteria combinations and showed the highest ratio values in prediction using Condition 3, 3 days after the 20-day learning period. As a result, when using the frequency ratio prediction model that learns for 20 days and predicts 3 days later and the DVC selection algorithm using Condition 3 (3 days in a row, the difference between the frequency ratio of the current and previous day is ≥ 0, the predicted frequency is > 10%, and the measured value exceeds the predicted value), all variant-defining POS-NTs are identified for Delta and Omicron with the highest ratio (Eq. (1), Table 5).

**Table 5:**
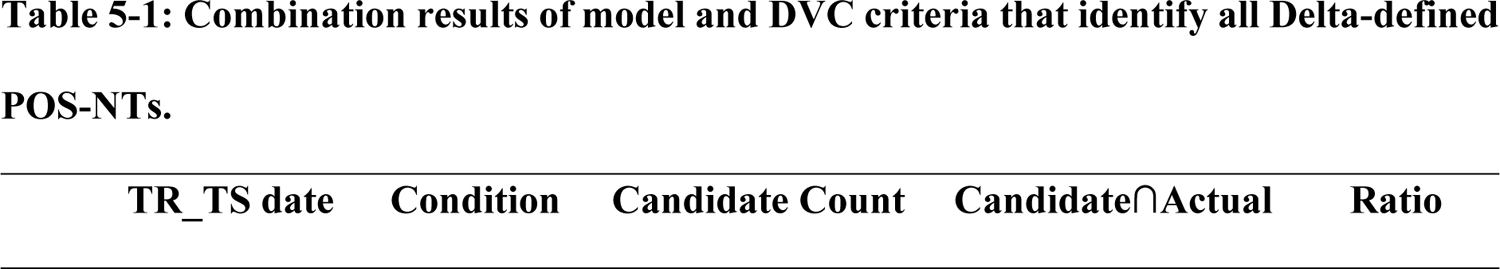
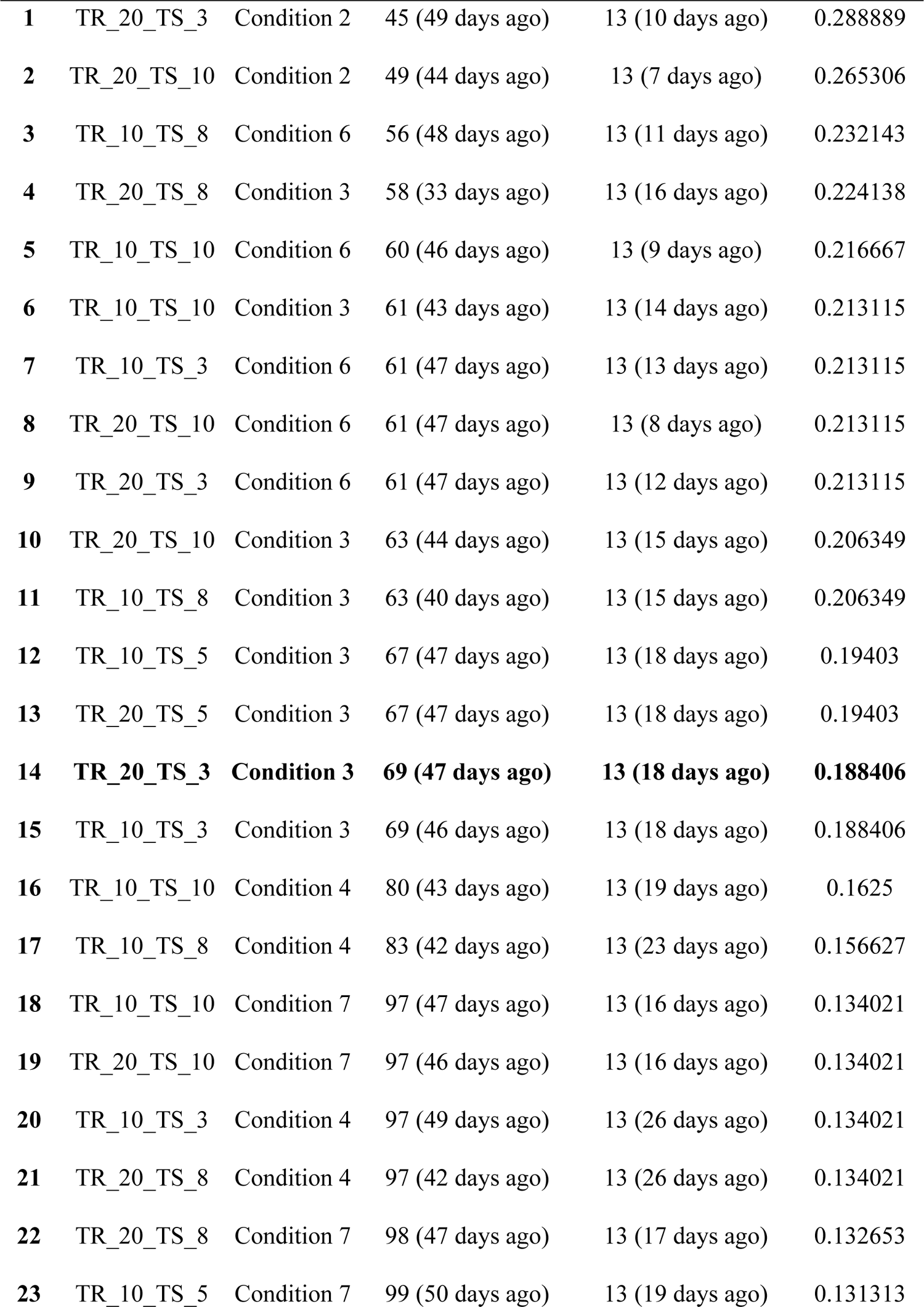
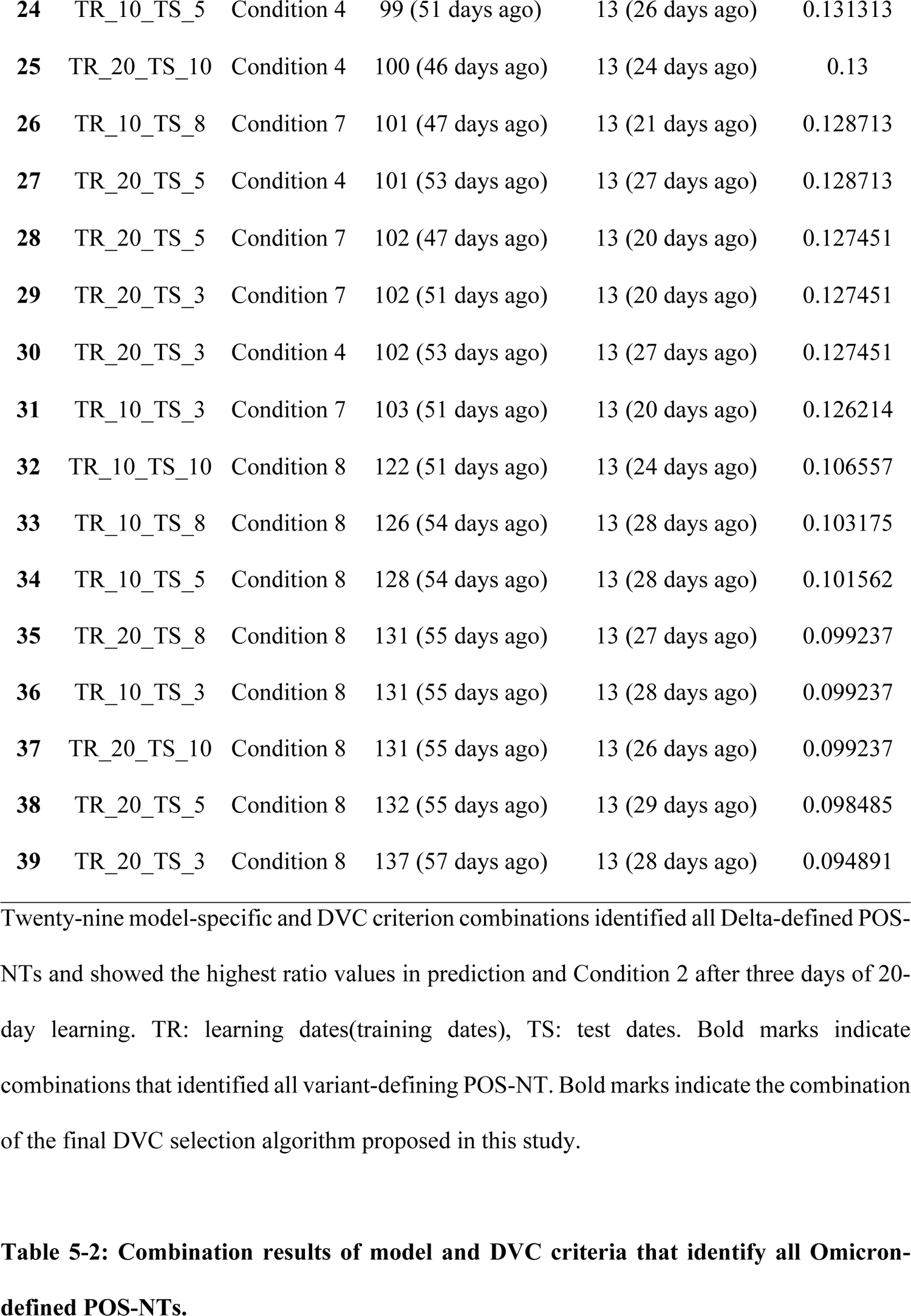
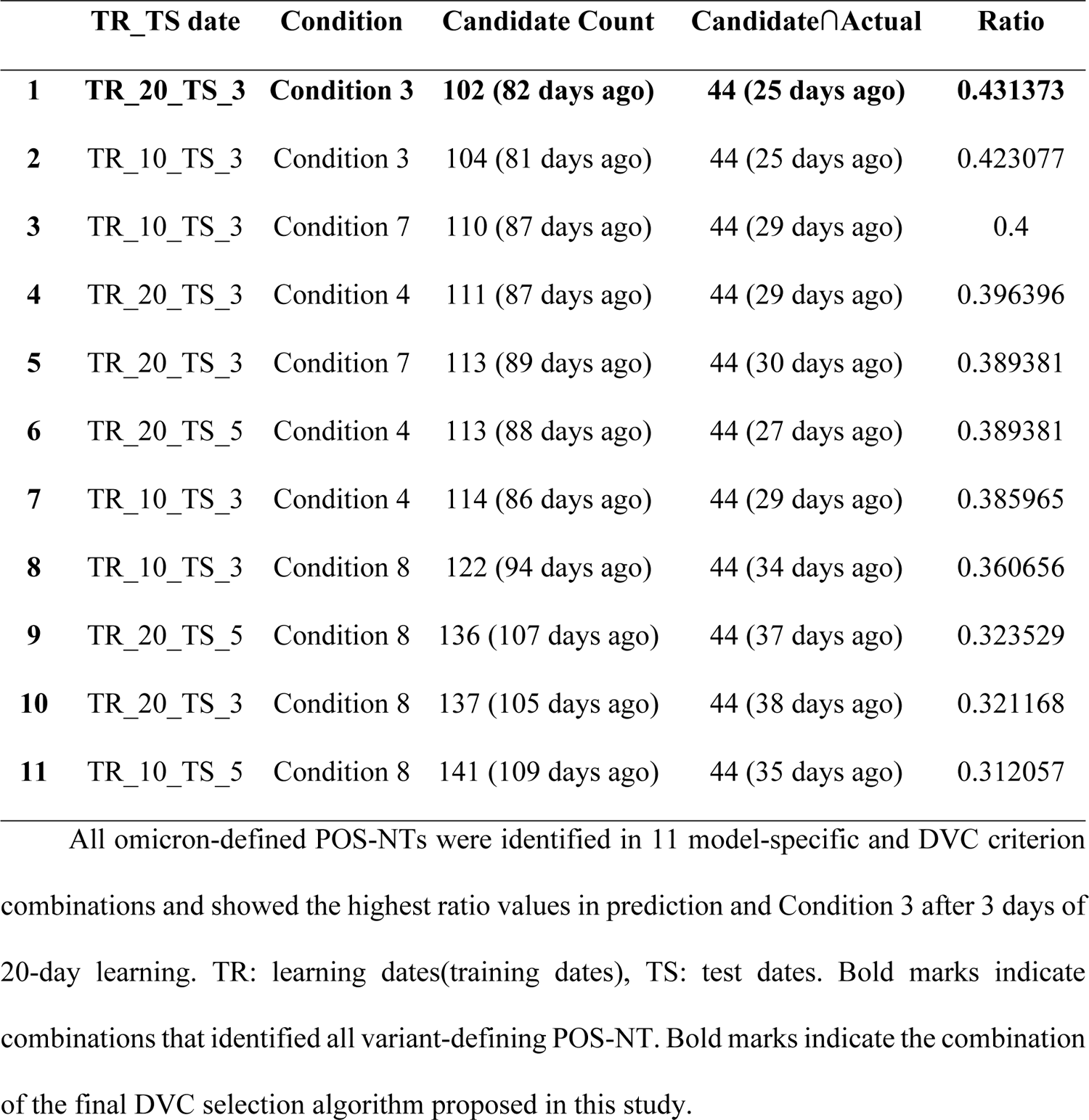
Combination results of POS-NT identification model and DVC criteria for all variant definitions.

Through the optimal ratio prediction model (i.e., learning for 20 days and prediction 3 days later) and DVC selection algorithm (i.e., Condition 3) 69 DVC POS-NTs were identified for Delta mutation, an average of 47 days before the dominant date. Among them, 13 Delta variant-defining POS-NTs were recognized 18 days before the dominant date. Similarly, 102 DVC POS-NTs were identified for Omicron mutation an average of 82 days before the dominant date, of which 44 Omicron variant-defining POS-NTs were recognized 25 days before the dominant date.

## Discussion

Many previous studies have predicted the incidence of COVID-19 and the ratio of Delta and Omicron mutations. For example, Pathan and Biswas predicted the COVID-19 time series by analyzing the ratio of 12 base mutations using 3,068 samples and the LSTM model from NCBI GenBank in 2020 to predict the mutation rate for future patients who do not yet exist [8]. Singh et al. obtained COVID-19 case count data for 15 states in India through the Kaggle website and predicted the future spread of SARS-CoV-2 using the Kalman filter [9]. Marzouk et al. collected the COVID-19 data of Engypt from the Flevy open source in 2021 and predicted COVID-19 outbreak (i.e., cumulative infection) after one week and one month, using LSTM, CNN, and MLP; the prediction results were in excellent agreement with the reported results [10]. Meanwhile, Obermeyer et al. proceeded with clustering using GISAID data on January 20, 2022, and the Pango lineage to infer prevalence for each lineage. Subsequently, they developed a hierarchical Bayesian regression model, PyR0, to detect and predict increases in B.1.1.7, AY.4, and BA.I in England [11]. De Hoffer et al. used 646.697 spike protein sequence data from the UK through GISAID in 2022 to perform clustering on a monthly or weekly basis based on amino acid substitution information and defined the appearance of a major cluster. They defined a new permanent variant as a chain containing clusters that share the same variant three or more consecutive times and designated an early warning for the emergence of a new permanent variant when 1% of the total sequence data was reached. As a result, an early warning was provided for the Alpha cluster as a new permanent variant six weeks before the WHO officially classified it as a VoC [12].

Although a few studies have predicted the occurrence of new mutations [Jankowiak, 12], they used protein-based data, and no studies have confirmed the trend by predicting the POS- NT ratio. Therefore, the current study can provide more detailed information regarding SARS-CoV-2 variants by predicting the trend and aspect of the mutation for each POS-NT.

This study has several limitations. First, the increasing POS-NT ratio was predicted using the DVC candidate selection algorithm, while the decreasing POS-NT ratio remained unanalyzed. Second, given that the dominant variant candidate identification algorithm was developed based on USA data, the algorithm may not apply to other countries in Asia. Hence, as different countries have demonstrated different rates of SARS-CoV-2 transmission and emergence of dominant variants, it is necessary to develop DVC selection algorithms for other countries, such as the UK, Germany, and Korea in the future. Third, only replacement mutations were analyzed in this study, whereas other mutation types, such as insertions and deletions, were not considered.

## Conclusions

We obtained SARS-CoV-2 POS-NT frequency ratio data for each country using a large amount of GISAID sequence data and defined the time point of the dominant variants for each mutation in each country. Subsequently, we developed a SARS-CoV-2 POS-NT frequency ratio prediction model and DVC selection algorithm using GPR for the USA and verified them for Delta and Omicron. Using this algorithm, we successfully identified all DVC POS-NTs before the dominant date, regardless of the soaring or gently increasing POS-NT patterns. As we were able to identify all mutation definitions of POS-NT for Delta and Omicron mutations, the algorithm can provide early warnings for other mutations in the future. If sufficient data exists, our model is expected to serve as an early warning algorithm for other viruses, thus, improving global health.

### Availability of the data and materials

The COVID-19 nucleotide sequence data used in this study can be obtained through GISAID (https://gisaid.org/) and compared with the original nucleotide sequence NC_045512. Correspondence and requests for materials should be addressed to TaeJin Ahn.

## Supporting information

**S1 Fig.**

**S2 Fig.**

**S3 Fig.**

**S4 Fig.**

**S5 Fig.**

**S6 Fig.**

**S7 Fig.**

**S8 Fig.**

**S9 Fig.**

**S10 Fig.**

**S11 Fig.**

**S12 Fig.**

**S13 Fig.**

**S14 Fig.**

**S1 Table.**

**S2 Table.**

## References

1. Rasmussen CE. Gaussian Processes in Machine Learning. In: Bousquet O, von Luxburg U, Rätsch G, editors. Summer School on Machine Learning. Springer; 2004. pp. 63–71.

2. Schulz E, Speekenbrink M, Krause A. A tutorial on Gaussian process regression: Modelling, exploring, and exploiting functions. J Math Psychol. 2018;85: 1–16.

3. Jarndal A, Husain S, Zaatar O, Al Gumaei T, Hamadeh A. In: 2020 International Conference on Communications, Computing, Cybersecurity, and Informatics (CCCI); 2020. pp. 1–5.

4. Tolba H, Dkhili N, Nou J, Eynard J, Thil S, Grieu S. GHI forecasting using Gaussian process regression: Kernel study. IFAC-PapersOnLine 2019;52: 455–460.

5. Velásquez RMA, Lara J VM. Forecast and evaluation of COVID-19 spreading in USA with reduced-space Gaussian process regression. Chaos Solit Fractals 2020;136: 109924.

6. Dhamodharavadhani S, Rathipriya R. COVID-19 mortality rate prediction for India using statistical neural networks and Gaussian process regression model. Afr Health Sci. 2021;21: 194–206.

7. Lounis M, Khan FM. Predicting COVID-19 cases, deaths and recoveries using machine learning methods. Eng Appl Sci Lett. 2021;4: 43–49.

8. Pathan RK, Biswas M, Khandaker MU. Time series prediction of COVID-19 by mutation rate analysis using recurrent neural network-based LSTM model. Chaos Solit Fractals 2020;138: 110018.

9. Singh KK, Kumar S, Dixit P, Bajpai, MK. Kalman filter based short term prediction model for COVID-19 spread. Appl Intell. 2021;51: 2714–2726.

10. Marzouk M, Elshaboury N, Abdel-Latif A, Azab S. Deep learning model for forecasting COVID-19 outbreak in Egypt. Process Saf Environ Prot. 2021;153: 363–375.

11. Obermeyer F, Jankowiak M, Barkas N, Schafner SF, Pyle JD, Yurkovetskiy, et al. Analysis of 6.4 million SARS-CoV-2 genomes identifies mutations associated with fitness. Science 2022;376: 1327–1332.

12. de Hoffer A, Vatani S, Cot C, Cacciapaglia G, Chiusano ML, Cimarelli A, et al. Variant-driven early warning via unsupervised machine learning analysis of spike protein mutations for COVID-19. Sci Rep. 2022;12: 9275.

